# Redefining paradigms in the archaeal virus-host arms race

**DOI:** 10.1101/2025.04.20.649705

**Authors:** Laura Martínez-Alvarez, Xu Peng

## Abstract

Archaeal antiviral defense systems remain poorly characterized despite recent advances in understanding prokaryotic immunity. Here, we analyze 7,747 archaeal genomes, the largest and most diverse dataset to date, revealing a striking disparity in defense system prevalence and diversity compared to Bacteria. Nearly one-third of archaeal genomes lack known systems beyond CRISPR-Cas and restriction-modification (in contrast to only 2.2% bacterial genomes), and only 31% contain CRISPR-Cas systems, far below previous estimates. While many known defense systems appear restricted to Bacteria, several single-gene candidate systems (PDCs) are enriched in Archaea. Phylogenetic analyses suggest that PDC-S27, PDC-S70, and PDC-M05 likely originated in Archaea, representing rare archaeal contributions to the prokaryotic immune repertoire. Consistent with earlier studies, our findings support the existence of deep evolutionary links between archaeal and eukaryotic systems for argonautes and viperins. These analyses highlight both the underexplored nature and the evolutionary significance of archaeal immunity, calling for expanded efforts to uncover archaeal-specific systems and improve our understanding of immune evolution across domains of life.

---

Mobile genetic elements (MGEs) are major drivers of horizontal gene transfer (HGT). They contribute to microbial genetic diversity and evolutionary innovation by facilitating the spread of adaptive traits such as antibiotic resistance and virulence factors through mechanisms such as transformation, transduction, and conjugation^1,2^. Their interactions with hosts range from mutualistic to parasitic, often imposing significant fitness costs^3,4^.

To counteract the costs imposed by MGEs, prokaryotes have evolved diverse defense strategies, from receptor modifications to sophisticated defense systems that degrade or modify nucleic acids, arrest cell growth, or disrupt membranes^5–7^. Defense systems often cluster in genomic “defense islands”, frequently alongside MGEs^8^. These are regions prone to horizontal transfer, an arrangement that may facilitate their synergy and co-mobilization^9,10^. The reciprocal selective pressure between MGEs and host defenses drives an ongoing evolutionary arms race, resulting in rapid innovation of prokaryotic immunity^4^.

Over 300 antiviral defense systems have been identified to date, many through recent advances in computational approaches^11–15^. Although more than one-third of known systems also appear in Archaea^5^, studies of archaeal immunity have remained limited in scope. Until recently, analyses relied primarily on RefSeq genomes, where archaea comprised fewer than 2% of the data^16,17^. This limitation has led to assumptions that archaeal immune landscapes largely mirror those of bacteria, with CRISPR-Cas and restriction-modification (RM) systems being the primary known components.

Two recent studies expanded defense analyses in Archaea but focused primarily on the *Asgardarchaeota* phylum or on specific systems (viperins and argonautes)^18,19^. Only CRISPR-Cas diversity has been comprehensively mapped across the domain^20–22^. Broader evaluations of other systems, including classical ones like restriction-modification, are lacking.

Meanwhile, archaeal genome availability has surged, especially through the recovery of uncultured lineages via environmental sequencing^23,24^, making a comprehensive re-evaluation archaeal defenses timely. Understanding archaeal immunity is key to reconstructing the evolution of antiviral defense across living organisms, especially given the proposed archaeal ancestry of eukaryotes^25,26^ and evolutionary connections between bacterial and eukaryotic systems^27–29^.

In this study we analyze 7,747 archaeal genomes, the largest and most taxonomically diverse dataset to date, to assess the distribution, diversity and evolution of antiviral defense systems in Archaea.

## Results

### Database creation and evaluation of defense system identification tools

We curated a comprehensive database of prokaryotic and metagenomic genomes, including 7,747 archaeal and 40,000 bacterial genomes from publicly available sources (Supplementary Table 1)^30,31^. Bacterial genomes were randomly subsampled from a total of 394,933 genomes available in the Genome Taxonomy Database (GTDB)^30^, while all archaeal genomes were included. Most genomes, 97.9% of bacterial genomes and 97.2% archaeal, exceeded >90% completeness and had <5% contamination; the remainder met GTDB’s inclusion criteria of ≥50% completeness and <10% contamination^30^. The taxonomic composition of the archaeal genomes is shown in Supplementary Figure 1.

To characterize archaeal antiviral immunity and compare it to bacterial systems, we used Padloc^16^ and DefenseFinder^17^. In addition, we benchmarked CRISPR-Cas Typer^32^ for the prediction of CRISPR-Cas systems, and a homology-based approach based on the REBASE database^33^ for the detection of restriction-modification systems (see Methods; Supplementary Tables 2-6).

To benchmark Padloc and DefenseFinder, we excluded proteins from unpublished candidate defense systems in Padloc to ensure fair comparison^15^. DefenseFinder and Padloc identified 150 and 118 systems, respectively (Supplementary Table 7). Overall, 50.9% of bacterial and 31.2% of archaeal defense proteins were identified by both tools (Fig. 1A). However, each tool also identified unique proteins: Padloc detected 1.6X more unique bacterial and 4.7X more unique archaeal proteins than DefenseFinder, with 56% of archaeal defense proteins uniquely detected by Padloc.

**Figure 1.**
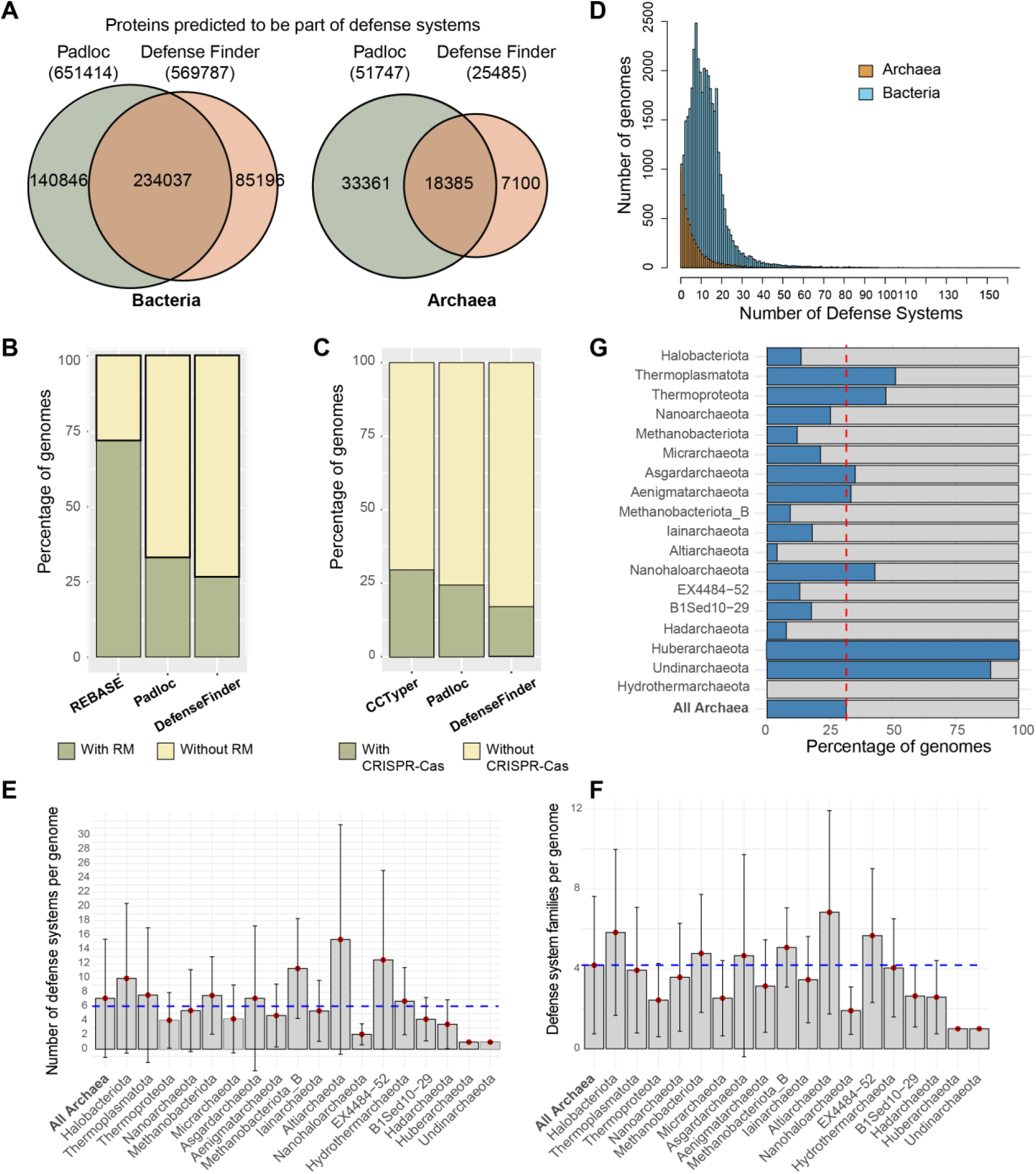
Benchmarking the identification of archaeal defense systems. **A)** Comparison of the output from Padloc and DefenseFinder on bacterial and archaeal datasets. **B-C)** Performance of tools in identifying restriction-modification systems **(B)** and CRISPR-Cas systems **(C)** within the archaeal dataset. **D)** Distribution of defense systems per genome across prokaryotic domains. The archaeal genome with the highest number of defense systems belongs to the *Thermoplasmata* class, with 71 systems. In bacteria, members of the *Polyangeaceae* family (*Myxococcota* phylum) have the highest count, with a maximum of 167 defense systems. **E-F)** Average number of defense systems **(E)** and defense system families **(F)** per genome across archaeal phyla **G)** Taxonomic distribution of archaeal genomes lacking defense systems other than RM and CRISPR-Cas, shown by phylum.

We further evaluated tool performance on RM and CRISPR-Cas system prediction. Padloc and DefenseFinder identified RM systems in 32.9% and 26.5% of archaeal genomes, respectively (Fig. 1B). These values are significantly lower than the estimated 81% prevalence reported in the REBASE database (709 archaeal genomes, October 2024)^33^. Both tools failed to detect many RM systems annotated in genomes listed in REBASE.

To improve detection, we developed a custom RM prediction pipeline based on REBASE. Archaeal proteins were matched to REBASE entries using MMseqs2 (≥65% identity, ≥80% coverage), and functional assignments were based on best hits. We further excluded proteins assigned to other defense systems by Padloc and DefenseFinder and classified RM system types based on gene neighborhood analysis (see Methods; Supplementary Table 6). This approach detected RM systems in 72.8% of archaeal genomes (Fig. 1B), consistent with previous estimates from smaller archaeal genome datasets^33–35^. Due to its higher sensitivity and agreement with prior benchmarks, we used this REBASE-based approach for all downstream analyses.

We also assessed CRISPR-Cas prevalence in Archaea using Padloc, DefenseFinder and CRISPR-Cas Typer. These tools identified CRISPR-Cas systems 24.5%, 16.9%, and 29.7% of archaeal genomes, respectively (Fig. 1C). The higher detection by CRISPR-Cas Typer was primarily due to putative II-D systems, which are known to be rare in Archaea^21,22^. Closer inspection revealed that many of these corresponded to OMEGA systems^36^, including IscB-HEARO^37^ (Supplementary Figure 2), rather than true type II-D CRISPR-Cas.

In summary, Padloc provided broader coverage (Fig. 1A) and superior detection of RM (Fig. 1B) and CRISPR-Cas (Fig. 1C) than DefenseFinder. Therefore, Padloc was selected for downstream analyses of defense system distribution.

### Domain-level overview of the prokaryotic defense landscape

We identified 521,796 occurrences of defense systems in bacterial genomes and 58,594 systems in archaeal genomes, comprising 1,049,452 genes in total (Supplementary Tables 2 and 6). This domain-wide analysis revealed clear differences in the distribution and diversity of defense systems between Archaea and Bacteria (Fig. 1D).

Defense system abundance deviated from a Gaussian “normal” distribution in both domains, but archaeal genomes showed a strong skew toward lower system counts per genome (Fig. 1D). Overall, 98.8% of bacterial genomes encoded at least one defense system, compared to 87.2% of archaeal genomes. Excluding RM and CRISPR-Cas, these proportions dropped to 97.84% and 68.37%, respectively (Fig. 1G), highlighting a more pronounced absence of known non-RM/CRISPR systems in Archaea.

In total, we detected 269 distinct system types in Bacteria and 197 in Archaea, with archaeal types representing 73.2% of all systems identified. Among these, only three system types were exclusively present in Archaea: CRISPR-Cas types III-G and IV-C, and a viperin system associated with a tetratricopeptide repeat-domain-containing protein, consistent with previous reports^38^. By contrast, 75 defense systems were exclusive to Bacteria, comprising 16,450 occurrences (3.15% of all bacterial hits), underscoring their relative rarity.

Bacterial genomes had an average of 14.4 defense system occurrences and 5.6 distinct types per genome, compared to 6.1 occurrences and 4.2 types in archaeal genomes (Fig. 1E-F; Supplementary Figure 3B). Defense systems were absent across a broad range of archaeal lineages (Fig. 1G), while their absence in Bacteria was rare (1.2%) and largely limited to taxa with reduced genomes and intracellular lifestyles (Supplementary Figure 3A), as reported previously^17^.

We observed a moderate positive correlation between genome size and defense system abundance in Bacteria (Spearman ρ=0.476, *p-*value < 2.2×10^-6^) and a weaker but significant correlation in Archaea (Spearman ρ=0.388, *p* -value = 1.169×10^-8^) (Supplementary Figure 3C).

The average genome size was 3.8 Mb for bacteria and 1.8 Mb for archaea, and many archaeal genomes belong to DPANN lineages (acronym for *Diapherotrites*, *Parvarchaeota*, *Aenigmarchaeota*, *Nanohaloarchaeota* and *Nanoarchaeota*), organisms with small genomes and cells^39^ (Supplementary Figure 1). Although genome size partially explains the reduced defense content in Archaea, these findings suggest that other lineage-specific or ecological factors also shape defense system distribution across domains.

### Archaeal defense systems show different prevalence than bacterial ones

To compare the immune landscapes of Archaea and Bacteria, we identified the 20 most prevalent defense systems in each domain. This analysis revealed 28 systems in total, representing the most widespread components of the prokaryotic immune repertoire, hereafter referred to as the core defensome (Fig. 2).

**Figure 2.**
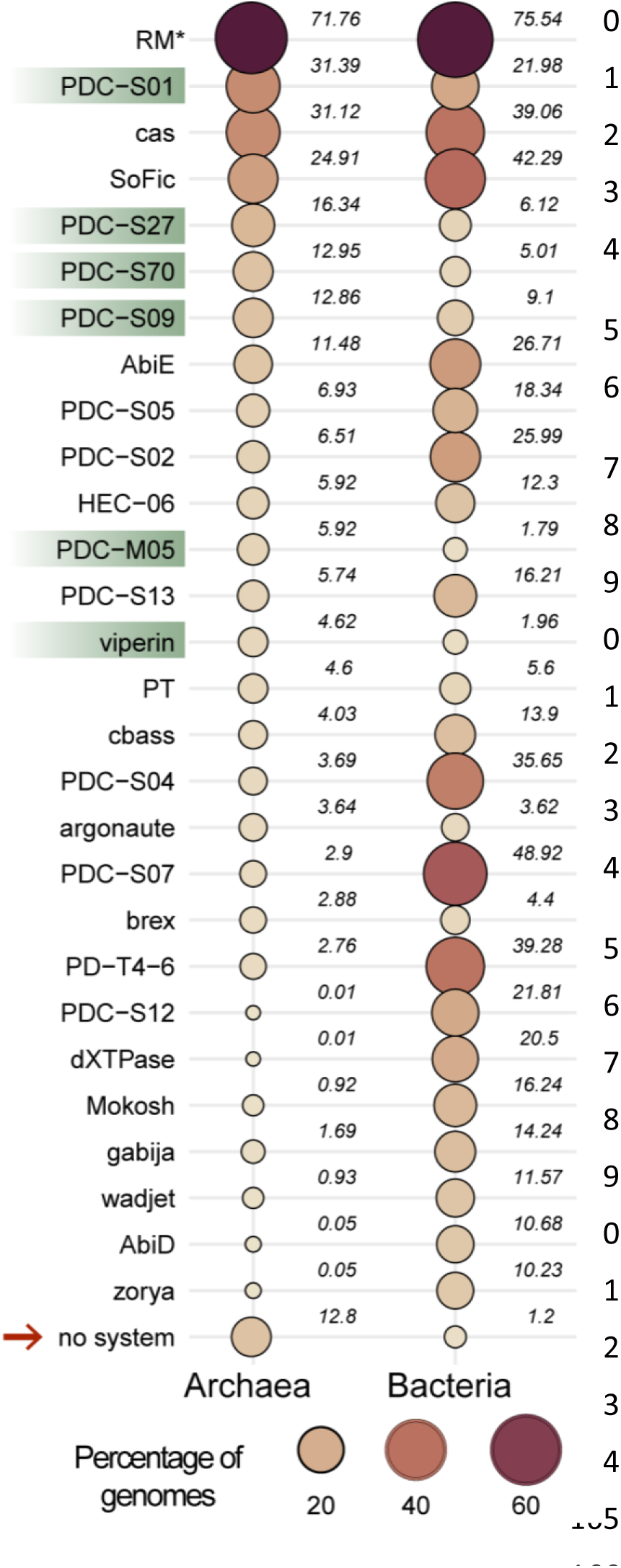
Prevalence of the 20 most abundant defense systems in Archaea and Bacteria. The bubble plot illustrates the percentage of genomes containing each defense system, with the size and color of circles indicating prevalence. Numerical values representing prevalence are shown to the right of each circle. Defense systems highlighted in green have higher prevalence in Archaea. The arrow indicates the percentage of genomes that lack any defense systems. Prevalence values are based on PADLOC outputs, except for archaeal RM systems (*), where prevalence was calculated using the REBASE-based approach (Methods and Fig. 1B)

Restriction-modification (RM) systems were the most abundant in both domains, found in 71.76% in archaeal and 75.54% bacterial genomes (Fig. 2). Despite similar prevalence, RM systems account for a disproportionately large share of the archaeal defense repertoire: 38.3% of all archaeal defense proteins, compared to 13.8% in Bacteria (Supplementary Figure 4). This difference is not due to greater RM copy number per genomes, as Archaea and Bacteria harbor comparable averages (2.0 vs 2.9 per genome). Instead, the elevated proportion in Archaea likely reflects the absence of many other defense protein homologs, increasing RM’s relative weight in archaeal immunity.

CRISPR-Cas systems were present in 31.1% of archaeal genomes and 39% of bacterial genomes (Fig. 2), contrasting sharply with earlier reports of 75-85% prevalence in Archaea^21,22^. These previous estimates were based on ∼300 genomes, heavily skewed toward hypehalophilic (60%) and hyperthermophilic (20%) species^40^. In contrast, our analysis, based on a 25 times larger dataset with broader taxonomic representation, likely provides a more accurate estimate of CRISPR-Cas prevalence in Archaea. Our results closely align with earlier bacterial estimates of 36-42%^21,22^.

Interestingly, six of the ten most prevalent systems are putative defense candidates (PDCs) (Fig. 2), recently identified single-gene systems discovered through a “guilt-by-embedding” approach^15^. Several Hma-embedded candidates (HECs), including HEC-06, have demonstrated antiviral activity in experimental assays^15^. Notably, PDC-S01 is the fourth most common defense across all prokaryotes and five PDCs (PDC-S01, PDC-S27, PDC-S70, PDC-S09, and PDC-M05) are more abundant in Archaea than in Bacteria (Fig. 2). These findings emphasize the relevance of PDCs to archaeal immunity and highlight them as promising targets for future characterization.

Among the 15 experimentally validated antiviral defense systems in the core defensome, only viperins are more prevalent in Archaea than in Bacteria, while argonautes show similar prevalence in both domains (Fig. 2). All other validated antiviral systems are more common in Bacteria, suggesting either greater evolutionary diversification in this domain or archaeal underrepresentation in current defense models.

The higher prevalence of most known antiviral systems in Bacteria, excluding RM, CRISPR-Cas, viperins, and argonautes, raises the possibility that these systems originated in Bacteria and were later acquired by Archaea through horizontal gene transfer.

### Lineage-dependent differences in the archaeal immune pangenome

We analyzed the distribution of experimentally validated antiviral systems with >3% prevalence across archaeal phyla, namely RM, CRISPR-Cas, SoFic, AbiE, viperin, DNA phosphorothioation, CBASS, and argonaute systems (Fig. 3). These systems show heterogeneous (“patchy”) distribution across archaeal lineages, a hallmark of defense system evolution likely shaped by frequent horizontal gene transfer^17^. A similar pattern is observed for PDCs, which have not been experimentally validated (Supplementary Figure 5).

**Figure 3.**
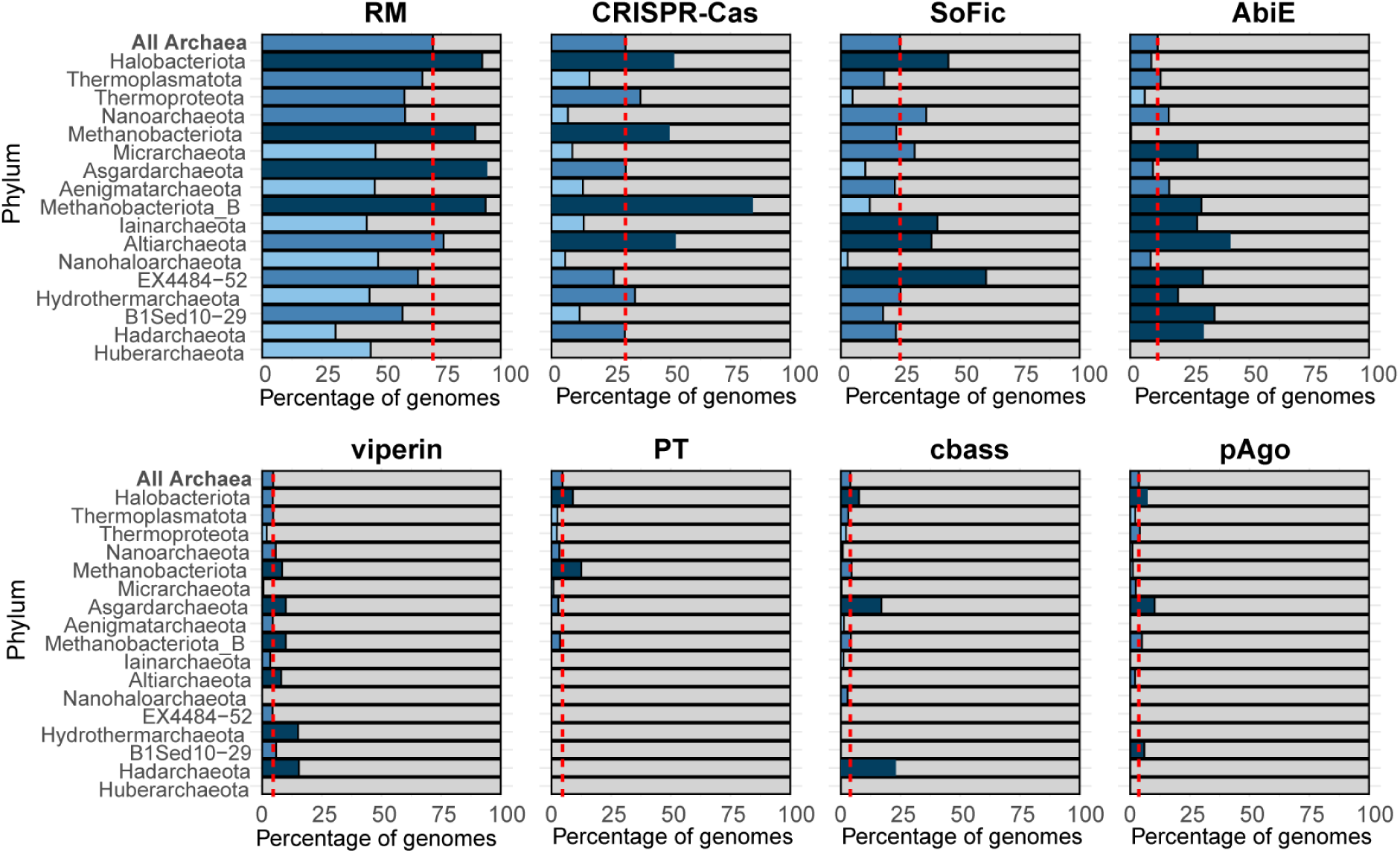
Taxonomic distribution of the archaeal core-defensome. The bar charts represent the percentage of genomes in each phylum containing the defense system. The red line indicates the average prevalence of each defense system across all archaeal genomes. Dark blue bars indicate overrepresentation of the defense system in a given lineage, while light blue bars reflect underrepresentation relative to the average.

Notably, DPANN archaea lack RM and CRISPR-Cas systems, yet show enrichment in AbiE SoFic and multiple PDCs. This mirrors patterns seen in host-dependent bacteria with small genomes (e.g. *Chlamydiota* and *Patescibacteria)*, suggesting a selective pressure for compact, single-gene systems in symbiotic or parasitic lineages (Supplementary Figures 3 and 5).

*Halobacteriota* members exhibit broad enrichment for both core and PDC defenses (Fig. 3, Supplementary Figure 5), with few underrepresented exceptions (e.g. Mokosh, PDC-S07). In contrast, *Thermoproteota* generally show lower defense prevalence, except for RM, CRISPR-Cas, and argonautes. *Thermoplasmatota* also exhibits low prevalence of CRISPR-Cas, but retains RM, AbiE, viperin, CBASS, and several PDCs (e.g. PDC-S01, PDC-S27, and PDC-S04). *Asgardarchaeota* genomes are enriched in RM, viperins, CBASS, argonautes and selected PDCs (S70, S09), but show underrepresentation of SoFic and most other PDCs (Fig. 3, Supplementary Figure 5).

We further examined the taxonomic and environmental distribution of CRISPR-Cas systems, given the ∼2.5-fold reduction in prevalence observed here compared to earlier estimates^21,22^. Prevalence varied across phyla and temperature classes. Although CRISPR-Cas prevalence was the highest in the hyperthermophilic *Methanobacteriota_B* phylum and in thermophilic classes of the *Thermoproteota* and *Halobacteriota* (>50% prevalence) than in mesophilic ones (10-25% and 0-37%, respectively), it was also enriched in mesophilic *Altiarchaeota* (∼50%) (Fig. 4A). DPANN archaea lack CRISPR-Cas entirely, regardless of their environmental temperature (Fig. 4A). While thermophily appeared to influence distribution, no significant correlation was found between CRISPR-Cas prevalence and predicted temperature (χ^2^(2) = 12.07, ρ-value = 0.0024). This suggests that temperature alone does not fully explain the distribution of CRISPR-Cas systems in Archaea.Ecological factors, particularly viral dynamics, may offer a more compelling explanation. Previous work has shown that environments with high viral abundance and low viral diversity are associated with higher CRISPR-Cas prevalence, whereas phylogenetic relatedness plays a limited role^41^. Symbiotic or parasitic lifestyles may further influence the selection of specific defense systems. Together, these patterns may help explain lineage-specific variation across archaeal clades.

**Figure 4.**
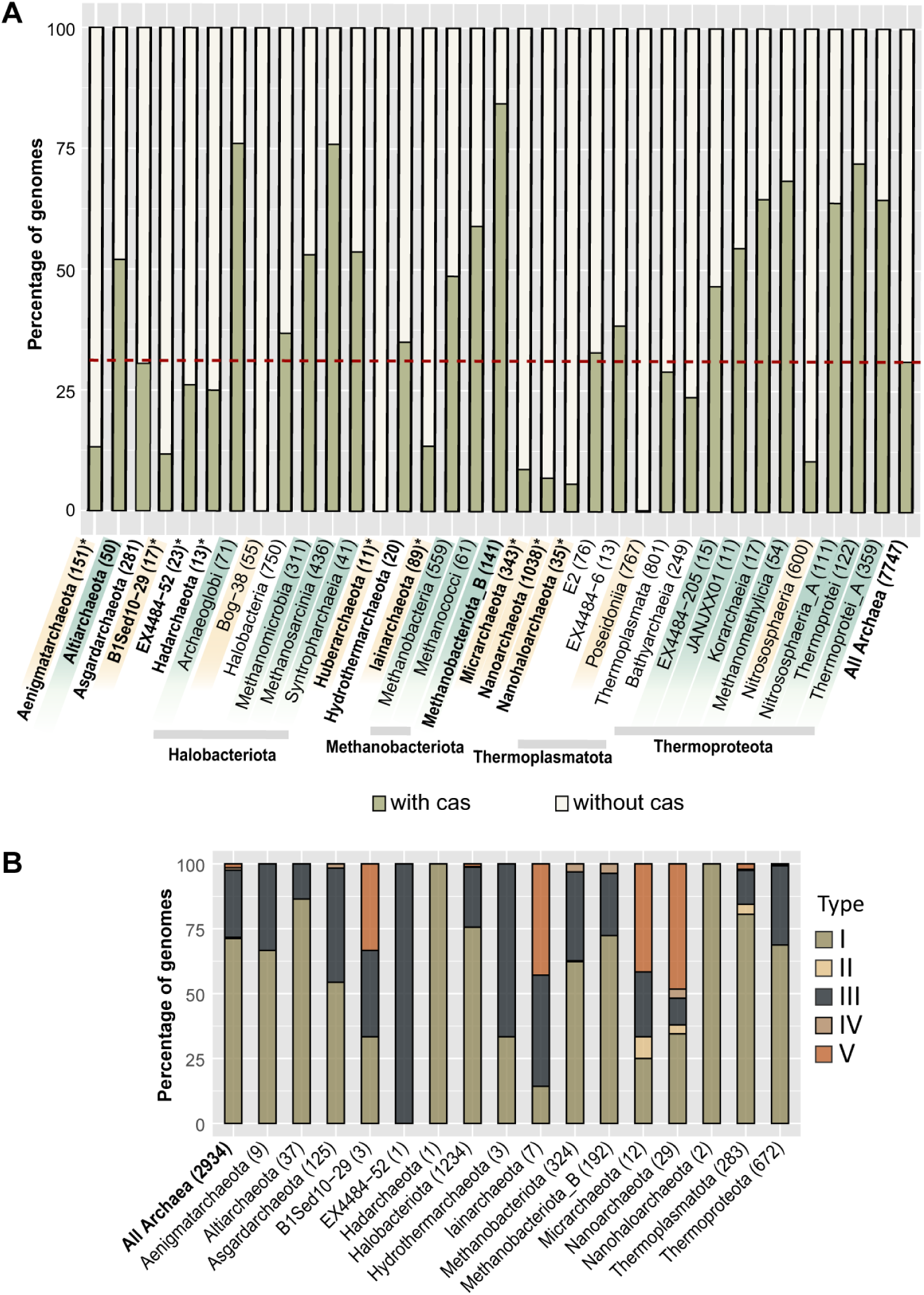
CRISPR-Cas distribution in Archaea. **A)** Prevalence of CRISPR-Cas system across archaeal lineages, with the red line indicating the average prevalence for the domain. Phylum names are in bold, while class-levels are shown in regular font. Lineages highlighted in green have above-average CRISPR-Cas prevalence, while those in yellow have below-average prevalence. Asterisk (*) denotes DPANN lineages. **B)** Relative abundance of CRISPR-Cas types across archaeal phyla. Numbers in parentheses indicate the total genomes analyzed for each lineage.

Despite reduced abundance, the distribution of CRISPR-Cas types remains consistent with prior studies: type I and III dominate, while type II and IV are rare, and type VI is absent (Fig. 4B)^21,22^. This suggests that, although prevalence varies, the retention of specific CRISPR-Cas types is governed by conserved evolutionary constraints.

### Evolutionary origins of the prokaryotic core-defensome

While the evolutionary trajectories of RM and CRISPR-Cas systems have been extensively reviewed elsewhere^20,22,34,35,42–46^, we focus here on the remaining components of the prokaryotic immune repertoire.

Several innate components in eukaryotes are linked to prokaryotic systems^27^. While many likely originated in Bacteria, others, such as viperins and argonautes, appear to have archaeal origins, particularly in *Asgardarchaeota*, the closest relatives of eukaryotes^18^. Although especially prevalent in *Asgardarchaeota*, these systems also appear across other archaeal phyla (Fig. 3).

To explore whether the current distribution of core defense systems reflects vertical inheritance or horizontal gene transfer (HGT), we analyzed the phylogenies of archaeal and bacterial homologs. In agreement with previous studies^38,47–49^, our trees support archaeal viperins and argonautes as ancestral immune systems, with deep phylogenetic roots (Fig. 5B,D). Both systems show patchy distribution with multiple inter-domain HGT events, but they are notably overrepresented in *Asgardarchaeota*, which make up only 3.6% of the archaeal dataset but account for 7.4% of archaeal viperins and 16% of archaeal argonautes (Fig. 3).

**Figure 5.**
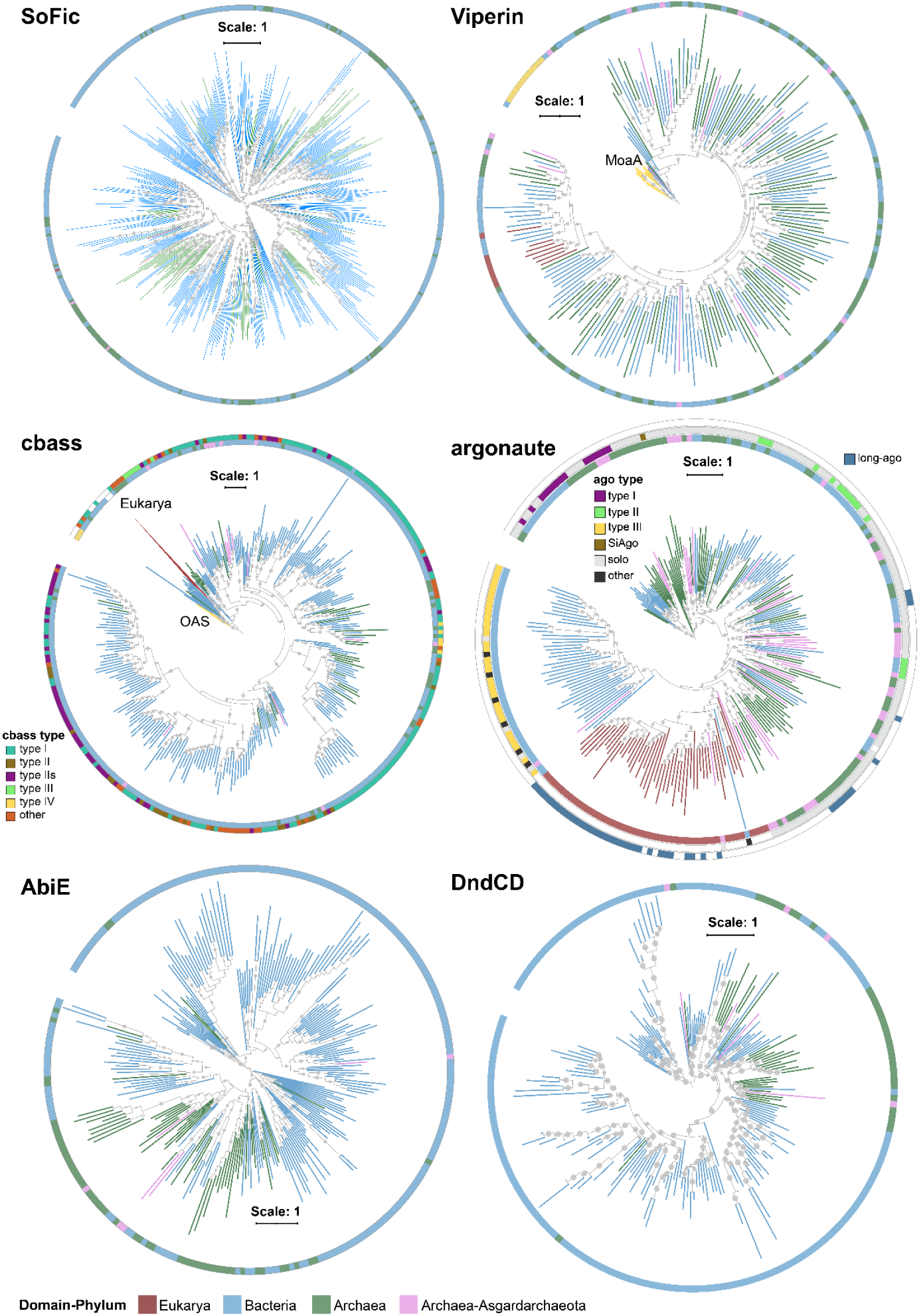
Evolutionary origins of the core-defensome. Phylogenetic trees of systems in the archaeal core-defensome. Branch and innermost ring colors indicate taxonomic classification: blue for bacteria, red for eukaryotes, green for archaea, and pink for asgard archaea. The outermost ring shows the system type classification. Bootstrap support values (≥70%) are marked as dots at the corresponding nodes. Trees for viperins **(B)** and CBASS **(C)** were rooted using MoaA and 2’-5’-oligoadenylate synthetase (OAS) sequences, respectively, while other trees were midpoint-rooted **(A,D-F)**.

Homologs of both systems are also found in eukaryotes^38,48^ and our analysis supports an archaeal origin for eukaryotic argonautes (eAgo), consistent with previous work^47,48^. Most asgardarchaeal argonautes fall within the clade from which eAgos appear to have originated (Fig. 5D), agreeing with previous work^18^. Similarly, the largest eukaryotic viperin cluster forms a sister group to a clade of archaeal proteins, including most asgardarchaeal viperins (Fig. 5B and Supplementary Figure 6). While this supports a likely archaeal origin for eukaryotic viperins, our analysis does not pinpoint a specific contributing phylum.

Our results also align with those of Culbertson and Levin^49^, who identified several bacterial-to-eukaryote HGT events involving viperins. However, in contrast to Shomar et al.^19^, who reported clear phylogenetic separation between archaeal and bacterial viperins, we observed no strict domain-based division (Fig. 5B and Supplementary Figure 6).

In contrast, systems such as SoFic, CBASS, AbiE and the DNA phosphorothioation (Dnd) appear to have a bacterial origin, with phylogenies indicating multiple bacteria-to-archaea transfer events (Fig. 5 A,C-E). Notably, while archaeal AbiE systems share a bacterial ancestor, they show domain-specific in phylogenetic trees (Fig. 5D). AbiE is a type IV toxin-antitoxin system known to induce abortive infection by acting on an unknown cellular target^50,51^, and its apparent domains restriction may reflect differences in host-specific targets.

PDCs display diverse evolutionary histories. Systems such as S09, S05, S02, S13, S04, S07, S12 and HEC-06 are predominantly bacterial and appear to have entered Archaea via HGT (Supplementary Figure 7). Others, like S01, S27, S70 and M05 are notably enriched in Archaea, suggesting an ancient archaeal origin (Supplementary Figure 7).

Among these, PDC-S01 and PDC-S27 are especially abundant in Archaea, accounting for 25% and 43% of their total detected occurrences. This corresponds to 9,986 S01 and 3,503 S27 systems in Archaea vs. 29,494 and 4,562, respectively, in Bacteria. Although S01 likely originated in Bacteria, it has undergone multiple transfers into Archaea, followed by domain-specific diversification (Supplementary Figure 7). Conversely, S27 appears to have originated in Archaea and spread into Bacteria (Supplementary Figure 7). Structurally, both encode ATPase domains fused to members of the PDDEXK superfamily, associated with nucleic acid targeting^15^.

PDC-M05 is even more archaeal-enriched (Supplementary Figure 7), comprising 59% of all detected instances. Its architecture, composed of a nucleotidyltransferase and a HEPN-domain protein, is characteristic of known prokaryotic toxin-antitoxin modules^52^, although its defensive role remains to be confirmed. A related bacterial system has been described^53^, but its antiviral role has not been tested. Similarly, PDC-S70, which encodes a PIN nuclease domain is predominantly archaeal (56%) and likely originated in Archaea, with multiple subsequent transfers into Bacteria.

Although most PDCs lack experimental validation, their high representation in Archaea, domain-specific enrichment, and molecular features suggest they are functionally relevant and may represent archaeal contributions to the immune repertoire. These systems are strong candidates for future characterization.

## Discussion

Archaea remain the least studied domain of life, including their antiviral strategies. While CRISPR-Cas immunity has been characterized in detail^20,54^, the broader archaeal immune landscape has received little attention, limiting our understanding of the evolution of innate immunity across life.

Here, we present the most taxonomically comprehensive analysis of archaeal antiviral defenses to date, revealing a markedly lower prevalence and diversity of immune systems in Archaea compared to Bacteria. These patterns raise important questions about the evolution of immunity and point to a largely untapped reservoir of archaeal defense systems. While such differences may reflect real biological and ecological variation, they also underscore long-standing methodological biases in how defense systems have been identified and studied.

Most detection approaches have been developed using large bacterial datasets, bacterial-centric protein profiles, and validated in models such as *E. coli*^11–13^. In contrast, archaeal genomes are underrepresented (only ∼2% of high-quality assemblies^30^), often fragmented, and typically originate from uncultivated or genetically intractable lineages^23,55–57^. These factors limit the identification and validation of archaeal immune systems.

Despite these challenges, our analysis reveals distinct features of archaeal immunity that challenge previous paradigms. Our CRISPR-Cas analysis (Fig. 2–3) illustrates how taxonomic bias can dramatically affect prevalence estimates, emphasizing the importance of comprehensive sampling. Notably, over half of the top-ranking defense systems in Archaea are single-gene putative defense candidates (PDCs), many of which are not associated with known defense islands^15^. This suggests that Archaea may encode immunity in noncanonical genomic contexts. Among these, three PDCs (S27, S70, and M05) are likely of archaeal origin, rare examples of archaeal-derived contributions to the prokaryotic immune repertoire.

These findings highlight the need for archaeal-centric and novel discovery strategies. Recent approaches such as regulatory motif mining in archaeal viruses^58^ and guilt-by-embedding analysis^15^ have already uncovered hundreds of new candidate anti-defense genes and defense systems. Moreover, exploring genomic regions enriched in mobile genetic elements (MGEs), beyond classical defense islands, has proven fruitful in both Bacteria^59–63^ and Archaea^64,65^, and will be essential for capturing the full diversity of archaeal immunity. The discovery of viral inhibitors of archaeal defenses remains limited^66–70^, further underscoring the need to expand beyond conventional approaches.

Our evolutionary analyses reinforce the view that eukaryotic immunity is a mosaic, shaped by both bacterial and archaeal inputs. Viperins and argonautes, which have clear eukaryotic homologs, appear to have originated in Archaea, highlighting the evolutionary relevance of archaeal immunity. Lastly, the relatively low biomass of Archaea, estimated at only 10% of that of Bacteria^71^, may limit the evolutionary space for defense innovation and help explain the prevalence of bacteria-to-archaea and bacteria-to-eukaryote horizontal gene transfers^27,49^.

In conclusion, while our work highlights a lower prevalence and diversity of defense systems in Archaea compared to Bacteria, this likely reflects methodological and ecological biases rather than a true lack of defenses. Ongoing advances in sequencing, computation, and experimental tools will accelerate the discovery of archaeal-specific systems, an exciting frontier for understanding microbial immunity, its evolution, and its biotechnological potential.

## Materials and Methods

### Data

The GTDB database (accessed on May 5, 2023) was used to retrieve the accessions and metadata for archaeal and bacterial genomes with over 50% completeness and < 20% contamination^24,30^. From this, we obtained a dataset of 7777 archaeal genomes and 40 000 bacterial genomes (the latter randomly subsampled from the 394 933 available bacterial entries), which were downloaded from NCBI^31^ to create the Archaea and Bacteria datasets, respectively (Supplementary Table 1). Proteomes for each genome were retrieved from NCBI when available, or predicted from the genomic sequence using Prodigal v2.6.3^72^. Supplementary Figure 1 depicts the taxonomic diversity of the genomes in the Archaea dataset.

### Identification of defense systems

To identify known defense systems in the archaeal and bacterial genomes we used DefenseFinder v1.2.2^17^, Padloc v2.0.0^16^ and CRISPRCasTyper v1.8.0^32^ with default settings. For identifying restriction-modification systems based on the REBASE repository (accessed on September, 2024)^33^, archaeal proteins were matched to the REBASE entries using MMseqs2 (release_15-6f452)^73^ with parameters set to --min-seq-id 0.65, --cov_mod 0 and –c 0.8. Proteins were assigned functions and type according to the best match based on e-value scores, resulting in a pool of candidate components for restriction-modification systems. To refine these candidates, we excluded proteins identified as part of non-RM systems by Padloc or DefenseFinder. An in-house script was then used to retrieve the genomic neighborhood of all proteins annotated as restriction enzymes, capturing five genes upstream and five genes downstream of each restriction nucleases. This neighborhood analysis allowed the identification of RM systems meeting specific criteria: Type I RM – presence of type I R,M and S components; Type II RM – presence of type II R and M components; Type III RM – presence of type III R and M components; Type IV – the presence of a type IV module alone; Type IIG – the presence of a type IIG module alone. A comprehensive list of predicted RM modules is available in Supplementary Table 6. Analyses and graphical visualization of data were carried out using R version 4.3.3 (2024-02-29)^74^, RStudio version 2024.04.1 Build 748^75^ and the package *ggplot2* v3.4.2^76^. The final comparison of the archaeal and bacterial immune landscapes was done using the output of Padloc after discarding entries labelled as DNA modification systems (DMS), which denote proteins involved in defense systems that modify DNA but cannot be classified as complete defense systems, and VSPR entries, which are not defense systems. For the identification of archaeal restriction-modification systems, the output of the REBASE-based approach described above was used instead of the PADLOC prediction.

### Statistical analysis

Statistical analysis about the prevalence, abundance and diversity of defense systems in the archaeal and bacterial datasets was done using the *vegan* package (v.4.1.3)^77^. The Shapiro-Wilk test showed evidence of non-normality for the genome size (W=0.893, *p*-value < 2.2 x 10^-6^ for archaea and W=0.968, *p*-value < 2.2 x 10^-6^ for bacteria), total system counts per genome (W=0.678, *p*-value < 2.2 x 10^-6^ for archaea and W=0.735, *p*-value < 2.2 x 10^-6^ for bacteria) and defense system diversity per genome (W=0.788, *p*-value < 2.2 x 10^-6^ for archaea and W=0.957, *p*-value < 2.2 x 10^-6^ for bacteria) distributions in the archaeal and bacterial datasets. Spearman’s rank correlation was used to assess the relationship between genome size and defense system abundance. A weak correlation was observed between genome size and system abundance in Archaea (ρ=0.388, *p*-value 1.169 x 10^-18^), as well as genome size and system diversity (ρ=0.376, *p*-value 4.38 x 10^-126^) in the archaeal dataset. In contrast, a strong positive relationship was observed between system abundance and system diversity (ρ=0.872, *p*-value < 2.2 x 10^-6^). In Bacteria, moderate correlations were observed between genome size and system abundance (ρ=0.476, *p*-value < 2.2 x 10^-6^) and diversity (ρ=0.580, *p*-value < 2.2 x 10^-6^), and a strong correlation between system abundance and system diversity (ρ=0.863, *p*-value < 2.2 x 10^-6^). The Kruskal-Wallis test did not show a statistically significant difference in cas abundance for thermophilic archaea (χ^2^(2) = 12.07, *p*-value = 0.0024) when genomes were classified according to their taxonomic affiliation at class level as “thermophilic” or “other”.

### Phylogenetic analyses

Components of defense systems identified by Padloc were used for all phylogenetic analyses, with for restriction-modification systems, which were analyzed using components of REBASE-based approach. For analysis of specific defense systems, concatenated DndC and DndD amino acid sequences were used for the DndABCDE phosphorothioation system, PIWI-domain components for argonautes, the AbiEii module for AbiE systems, and the cyclase module for CBASS. Eukaryotic sequences were obtained from the following sources: eukaryotic viperins from Shomar et al.^19^, cGAS-like pattern recognition receptors (cGLRs) from Li et al.^78^ and argonautes from Swarts et al.^48^.

To reduce sequence redundancy, we clustered the components of each defense system using cd-hit v4.8.1 at a 65% similarity threshold^79^. Amino acid sequences were then aligned with MAFFT v7.505 using the *“auto”* option^80^ and trimmed with TrimAl v1.5.rev0 ^81^ using the *“gappyout”* setting. Preliminary phylogenetic trees were constructed with FastTree v2.1.11^82^ with parameters *–lg* and *–boot 100*. The preliminary trees were pruned for retaining sequence diversity using Treemer v.0.3^83^, resulting in a reduced dataset of sequences for constructing the phylogenetic trees presented in Figure 5. For these trees, sequences were newly aligned using MAFFT with parameters *--maxiterate 1000* and *--localpair*, and then trimmed with TrimAl using the *-gt 0.3* option. Final phylogenetic trees were constructed with IQ-TREE v2.0.7^84^ using options *–m MFP –bb 1000 –alrt 1000* and *-bnni* to select the best-fit model using the Model Finder algorithm, ultrafast boot approximations and approximate likelihood ratio tests with one thousand replicates each to assess branch support. Phylogenetic trees were visualized using ITOL^85^. Trees were rooted at midpoint, except for viperin and CBASS trees, which were rooted using MoaA and OAS genes, respectively, as an outgroup, following previous analyses^19,49^. PDC-S27 tree was rooted using AAA-ATPases (PF00004) as the outgroup.

The amino acid sequences and phylogenetic trees used to make Fig. 5 and Supplementary Figure 7 are deposited in Supplementary Table 9 and Supplementary Data.

## Acknowledgements

We acknowledge the support of the EcoCluster initiative at the Department of Biology of the University of Copenhagen for access to resources used for bioinformatics analyses.

## Author contributions

L.M-A. conceived the paper, performed the analyses and wrote the manuscript. X.P. revised and approved the manuscript.

## Supplementary Figures

**Supplementary Figure 1.**
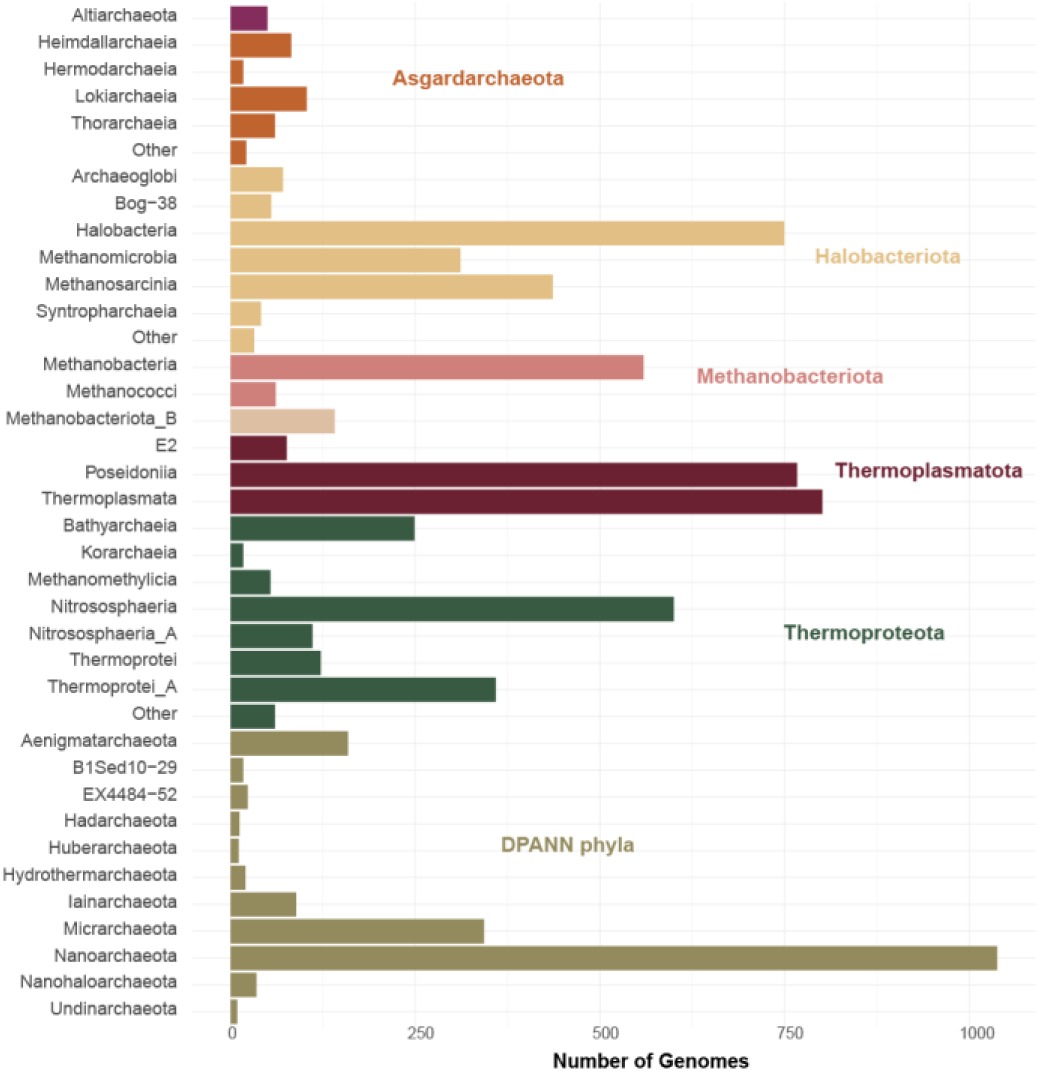
Taxonomic diversity of the archaeal database. The taxonomic classification of genomes follows the phylogeny of the Genome Taxonomy Database (Rinke et al. 2021). Bars are colored according to the phylum or superphylum to which they belong.

**Supplementary Figure 2.**
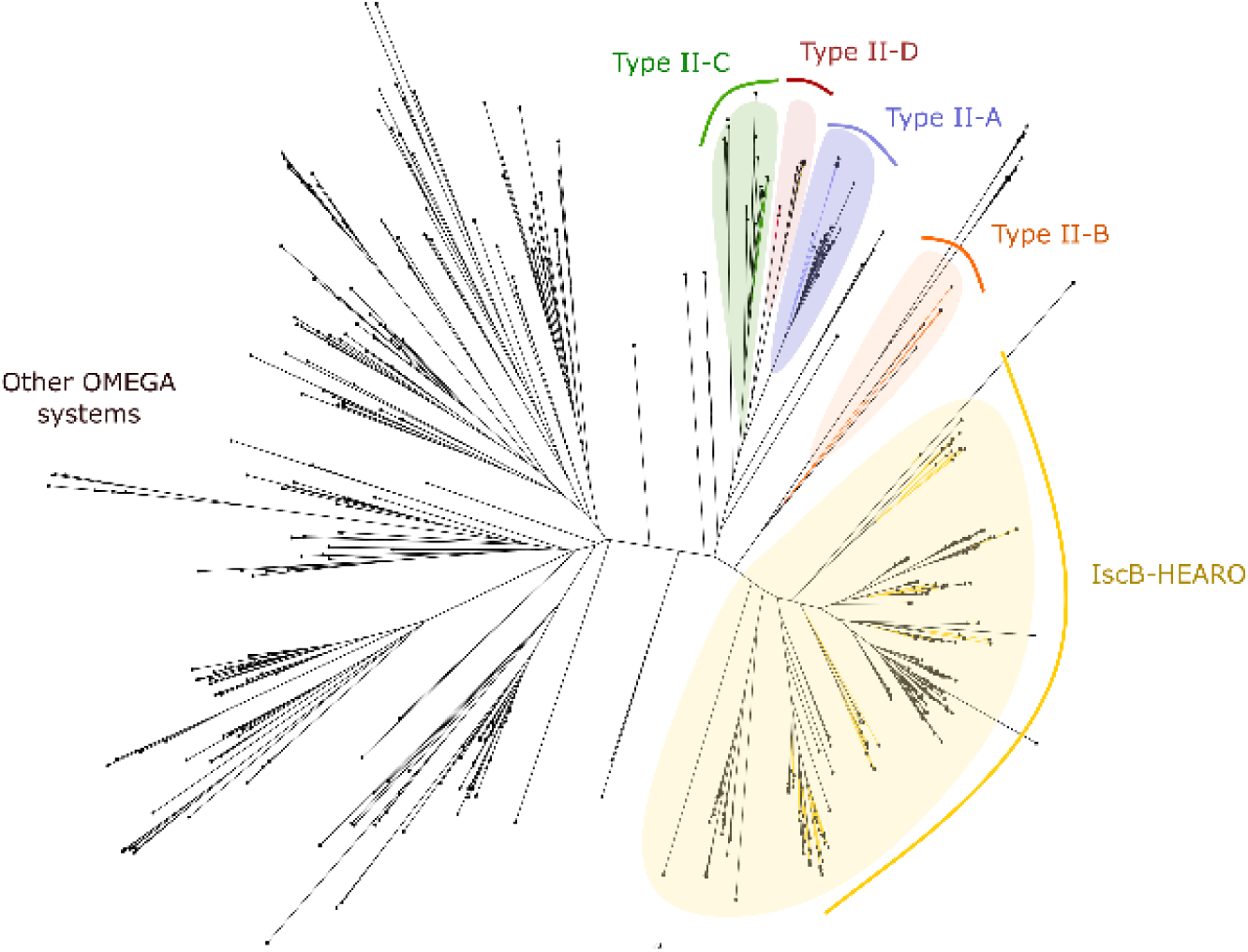
Phylogeny of archaeal type-II CRISPR-Cas systems identified by CRISPR-Cas Typer. Proteins from the archaeal database are indicated in black and representative sequences of the previously characterized Cas9 subtypes (II-A to II-D) or IscB elements are shown in color. Sequences belonging to OMEGA clades (Altae-Tran et al. 2021, Aliaga-Goltsman et al. 2022) are shaded in yellow. Proteins identified as type II-D CRISPR-Cas effectors do not cluster with the known Cas9 subtypes and are likely OMEGA systems.

**Supplementary Figure 3.**
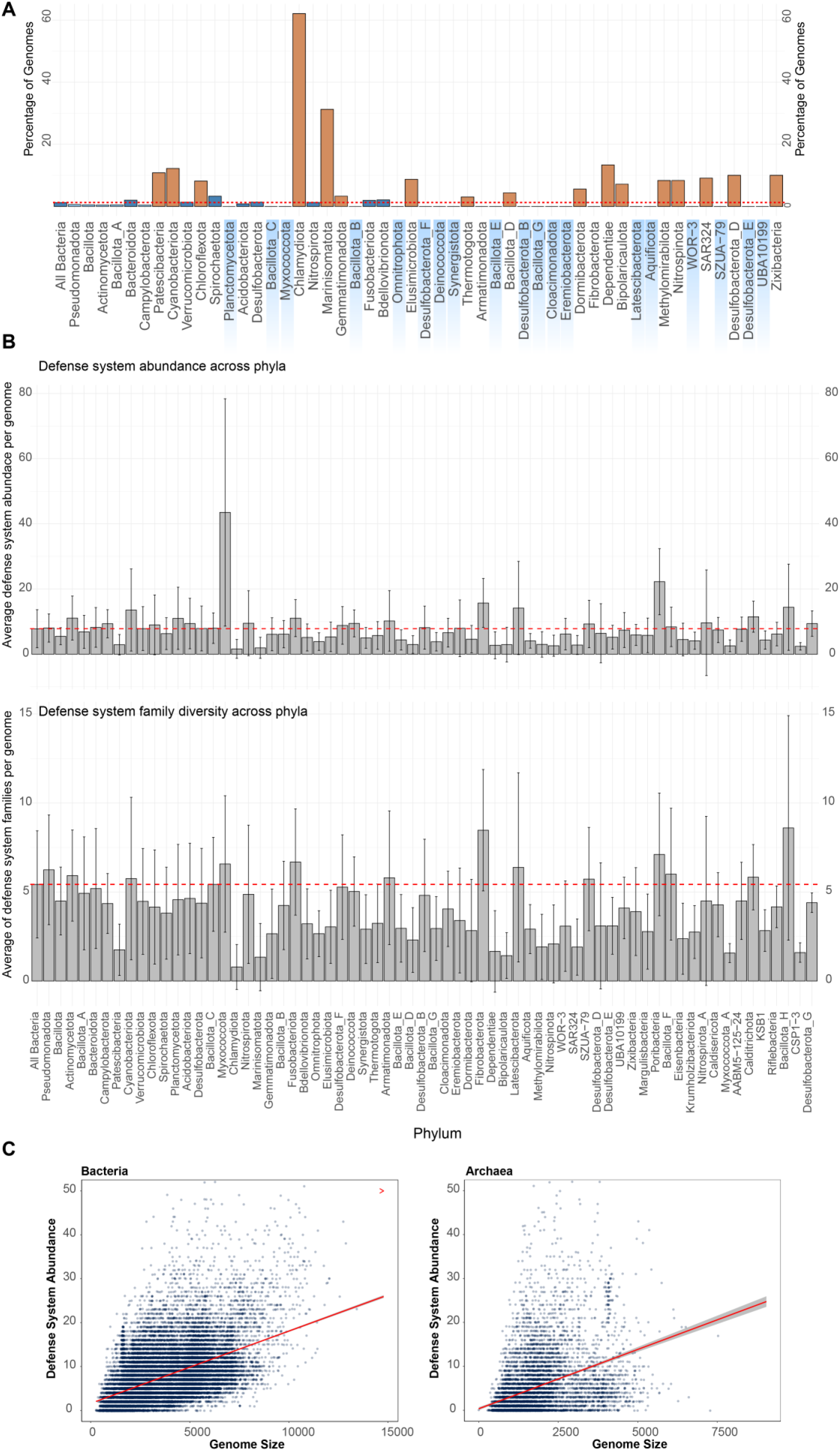
Statistical analysis of bacterial defense systems. **A)** Taxonomic distribution of bacterial genomes lacking defense systems. The dashed red line indicates the average percentage across all bacterial genomes analyzed. Orange bars indicate above-average values. All genomes belonging to the phyla shaded in blue encode one or more defense system. **B)** Comparison of defense system abundance and diversity across bacterial phyla. The top panel shows the average number of defense systems per genome. The bottom panel presents the average number of defense system families per genome. The dashed red line indicated the average value of all bacterial genomes analyzed. Error bars indicate standard deviation. **C)** Relationship between genome size and defense system abundance in Bacteria (Left) and Archaea (Right). Each dot represents a genome, with the number of defense systems plotted against genome size. The dotted red line represents the linear regression trendline, while the shaded area indicates the standard error of the regression fit.

**Supplementary Figure 4.**
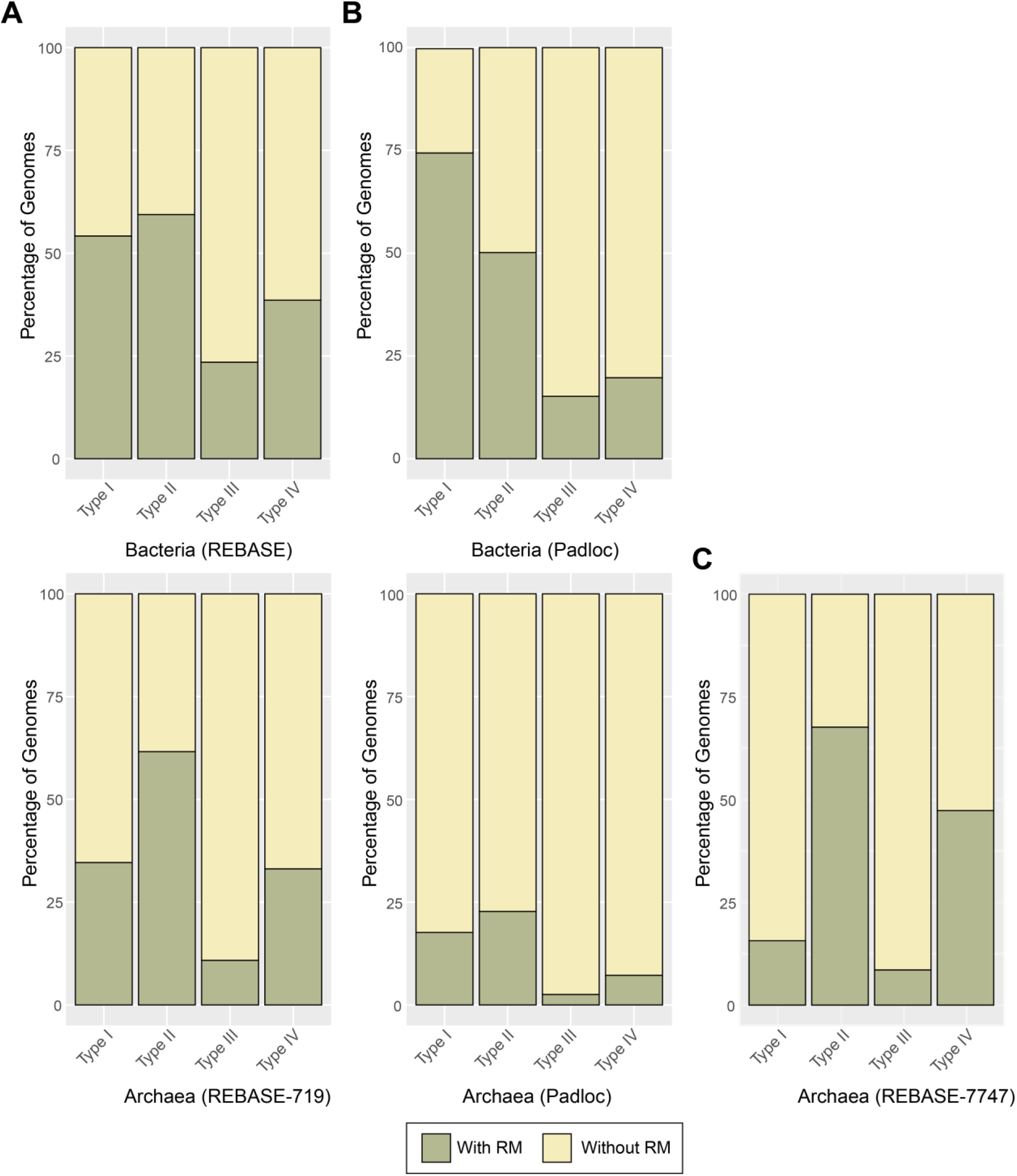
Output of the restriction-modification identification approaches. Prevalence of the four types of restriction-modification systems in the genomes deposited in REBASE (Roberts et al. 2023) **(A),** or in the bacterial and archaeal datasets used in this work as predicted by PADLOC **(B)** or our homology REBASE-based approach (REBASE-7747) **(C)**.

**Supplementary Figure 5.**
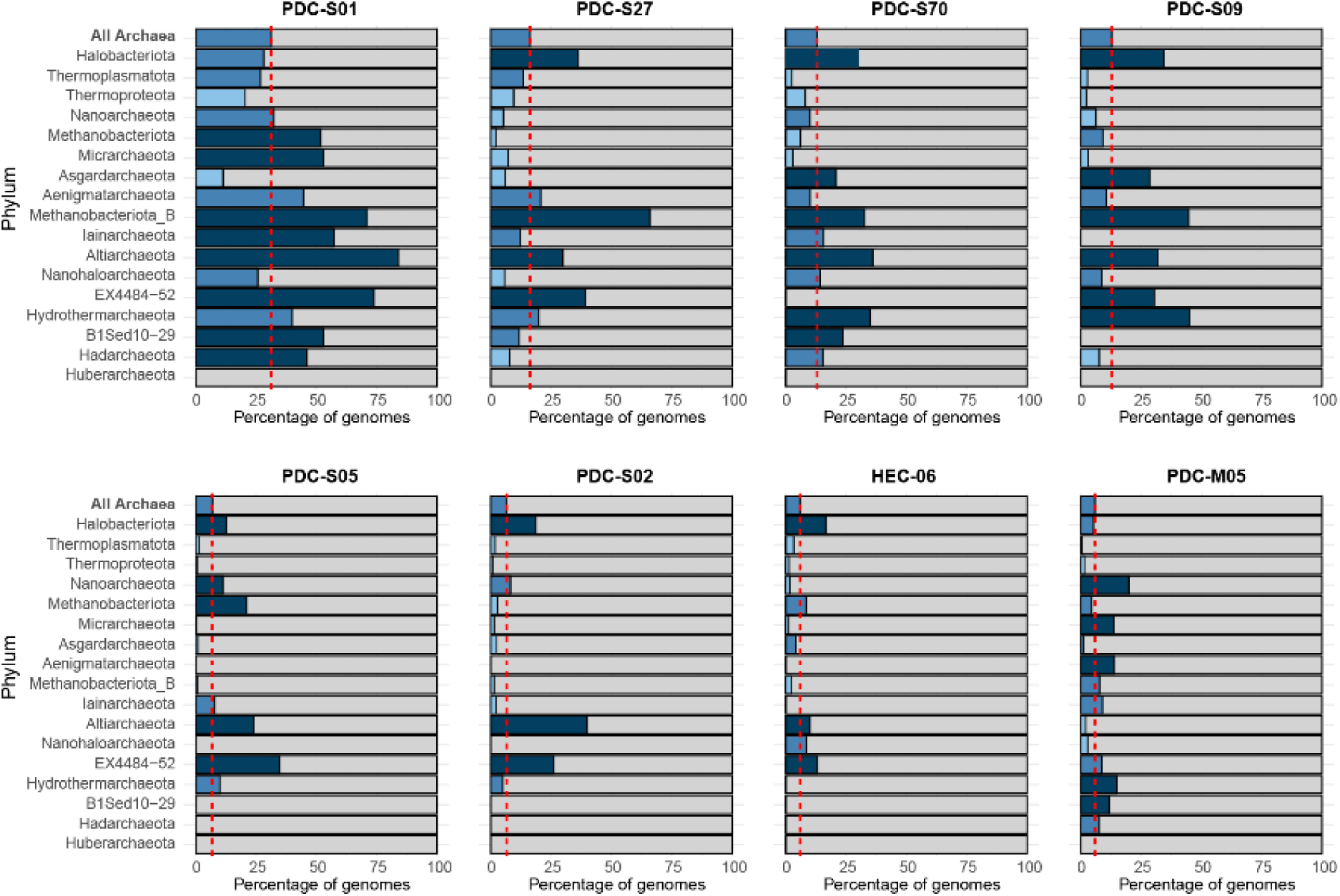
Taxonomic distribution of the archaeal PDC-core defensome. PDC systems in the top 20 most prevalent archaeal antiviral systems. The bar charts represent the percentage of genomes in each phylum containing the defense system. The red line indicates the average prevalence of each defense system across all archaeal genomes. Dark blue bars indicate overrepresentation of the defense system in a given lineage, while light blue bars reflect underrepresentation relative to the average.

**Supplementary Figure 6.**
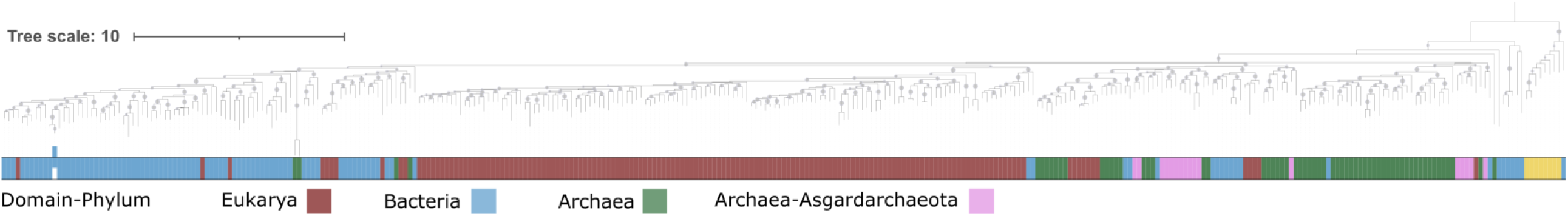
Phylogenetic analysis of viperins closely related to eukaryotic viperins. Prokaryotic viperins that clustered near eukaryotic viperins in a preliminary, broad phylogenetic tree (see Methods) were extracted and re-analyzed to investigate the evolutionary origins of eukaryotic viperins. The color strip indicates taxonomic classification: blue for bacteria, red for eukaryotes, green for archaea, and pink for asgard archaea. Bootstrap support values (≥70%) are marked as dots at the corresponding nodes. Tree was rooted using MoaA sequences (yellow).

**Supplementary Figure 7.**
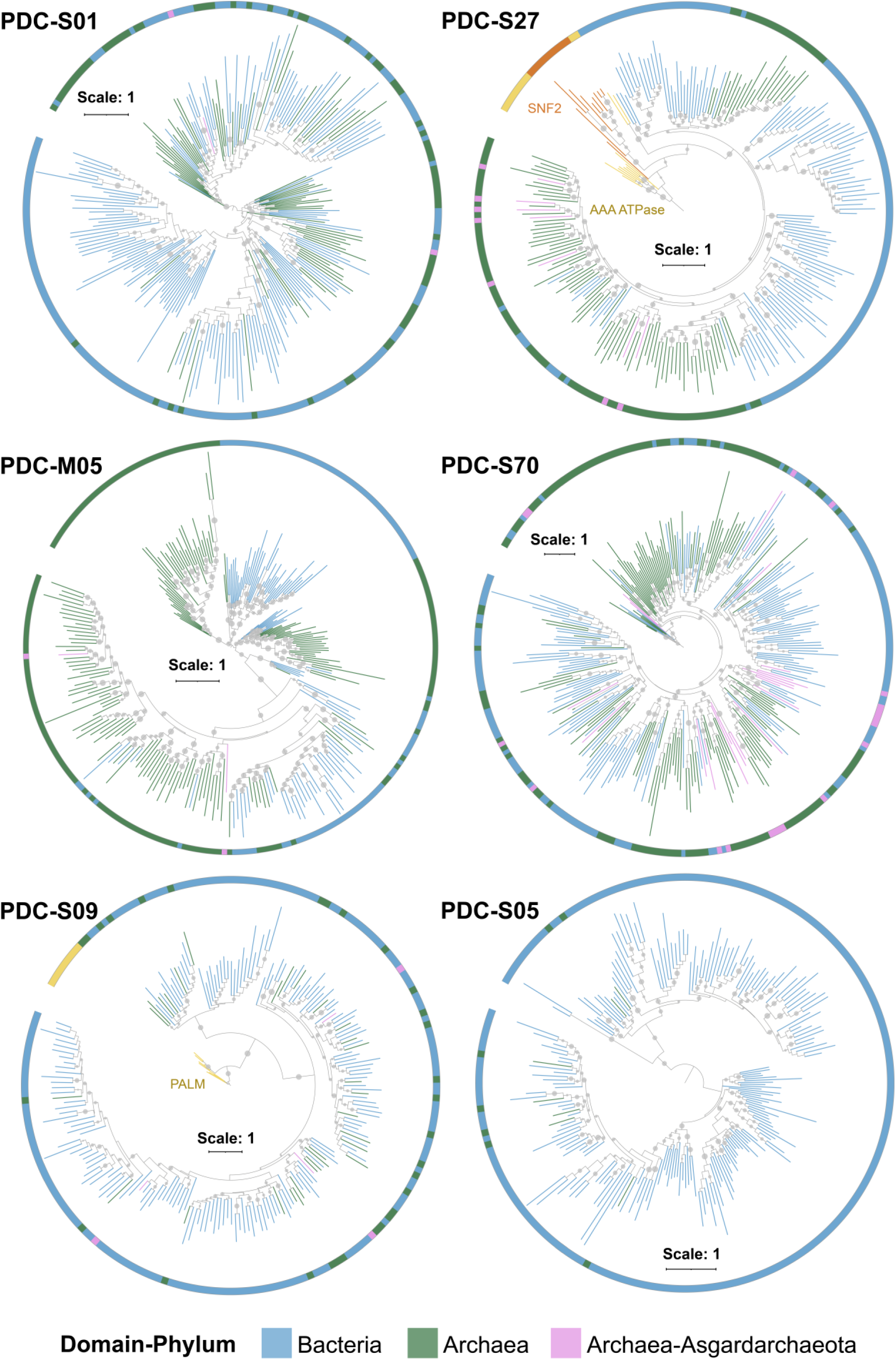
Evolutionary origins of the PDC-core defensome. Phylogenetic trees of the most prevalent PDC systems in the prokaryotic core-defensome. Branch and innermost ring colors indicate taxonomic classification: blue for bacteria, red for eukaryotes, green for archaea, and pink for Asgard archaea. Bootstrap support values (≥70%) are marked as dots at the corresponding nodes. Trees were rooted at midpoint, except PDC-S27 and PDC-S09, which were rooted using AAA ATPases (PF00004), and DNA polymerase beta PALM domain (PF14792) sequences, respectively (yellow).

